# Instilling Good Knowledge, Attitude and Practices among the Indigenous People of Malaysia Concerning Dog Associated Zoonotic Infections

**DOI:** 10.1101/381350

**Authors:** Abdul Rashid, Lau Seng Fong, Puteri Azaziah Megat Abd Rani, Siti Fatimah Kader Maideen, Intan Nur Fatiha Shafie, Nur Indah Ahmad, Farina Mustaffa Kamal, Mokrish Ajat, Sharina Omar

## Abstract

**Background:** The Jahai, a subethnic of the indigenous people of peninsular Malaysia, have commonly used dogs for hunting but have started to move away from traditional hunter-gatherer lifestyle, leaving dogs which were commonly used for hunting to wander around the villages and to multiply in numbers.

**Objective:** The objective of this study was to instil good knowledge, attitude and practices of the Jahai community concerning dog associated zoonotic infections using One Health concept.

**Methods:** This non-experimental pre and post-test intervention study was conducted among Jahai villagers aged 12 years and above living in a village located in the Belum forest in Malaysia. Interventions included health education and promotion using discussions, posters, slide presentations, comics and video clips with relevant content. In addition the children of the village were taught correct hand washing techniques and dog associated zoonotic infections.

**Results:** In general most aspects of knowledge, attitude and practice improved post intervention. The knowledge on risk of infections transmitted from pet dogs (X^2^=4.293, p= 0.038) and the practice of washing hands before eating (X^2^=14.984, p <0.001) improved significantly. The increase in the mean scores of the participants knowledge (t=−9.875, p=<0.001) and attitude (t= −4.100, p=<0.001) post intervention was statistically significant.

**Conclusion:** This study showed the effectiveness of a multidisciplinary team using One Health concept to successfully improve knowledge, attitude and practices related to dog associated infections. A sustained and committed health education and promotion interventions involving the community and school children in promoting heath should be custom made for indigenous communities, and sanitation and hygienic practices reinforced at every opportunity.

**Author’s Summary:** The indigenous people of peninsular Malaysia are a marginalized group; they are socio economically deprived and have low levels of education. One such group is the Jahai, who commonly used dogs for hunting, but have recently started to move away from traditional hunter-gatherer lifestyle, resulting in the dogs to multiply in numbers and roam as strays in the village. The community is now at risk of dog associated zoonotic infections. Studies have shown that health education and promotion can improve knowledge, attitude and practices of dog associated infections. However most of the studies were done among dog owners and in communities with fairly good education levels. A holistic approach using One Health concept was used to instil good knowledge, attitude and practices of the Jahai community concerning dog associated zoonotic infections. This non-experimental pre and post-test intervention study was conducted among Jahai villagers aged 12 years and above living in a village located in the Belum forest in Malaysia. The findings of this study showed the effectiveness of a multidisciplinary team using One Health concept to successfully improve knowledge, attitude and practices related to dog associated zoonotic infections.

## Introduction

It is estimated that there are more than 370 million indigenous people spread throughout 70 countries worldwide practicing unique customs and having distinct languages, social, cultural, economic, and political characteristics [1]. So much so that the United Nations issued a declaration on the rights of indigenous people to guide member states to form national policies in order to protect the collective rights of indigenous people including their culture, identity, language and access to employment, heath, education and natural resources. Unfortunately indigenous people are usually neglected segments of society, derelict of political participation, economically marginalized, lack access to social services and are often discriminated. Large proportions of the indigenous people live in and around forests and are mostly hunter-gatherers.

‘Orang Asli’ is the Malay term literally meaning ‘original people’ was used by the British to refer to the indigenous minority living in peninsular Malaysia. They are believed to have inhabited the peninsula for up to 25,000 years [2]. There are three major groups of the peninsular aborigines; Negrito, Senoi and Proto-Malay, each with many sub-ethnic groups. According to a 2010 data they number about 160,000. The Orang Asli, like most other aboriginal people, are marginalized with very little resources and laden with socio economic and health problems. The Orang Asli lag very far behind in basic infrastructure, literacy and education resulting in very high poverty rates [3]. Although in recent years the health services provided to the Orang Asli has improved but despite this the morbidity and mortality rates among them is still high.

Since time immemorial the Orang Asli who are mostly hunter and gatherers are nomadic or semi-nomadic communities living in the forests of Peninsula Malaysia [4]. The Jahai, a subethnic group of Negritos, mostly live in the Royal Belum State Park, situated in Perak, one of the 14 states in Malaysia. The park is 117,500 ha and borders Thailand [5] and houses an estimated 219 families [6]. The Jahai who commonly use dogs for hunting have begun to move away from traditional hunter-gatherer lifestyle, selecting for more modern agricultural and fishing practices [7] as well as working in the tourism industry which has mushroomed in the park to subsidise their livelihoods and subsistence. Dogs which were commonly used for hunting are now left to wander around the villages. In one village of about 400 people it is estimated that there were about 100 dogs most of which have no owners.

Dogs can transmit zoonotic infections to humans [8–10] and considering that even vaccinated pet dogs can transmit zoonotic infections [11–14], the risk from stray dogs is higher. Humans are at risk of zoonotic infections from dogs through bites, scratches, direct or indirect contact with dog saliva, urine and other body fluids or secretions and by ingestion of dog faecal material, inhalation of infected aerosol or droplets and through arthropods or other invertebrate vectors like fleas, mites, ticks and lice. The risk is higher in people with under-developed immune system such as in the very young (<5 years), elderly (>65 years), pregnant women, or immunocompromised persons [8,9,11–13,15–18]. Because of the stray dog population in the Orang Asli villages, they are at risks of dog related infections which include rabies, noroviruses, *Pasteurella, Salmonella, Brucella, Campylobacter*, leptospirosis, giardiasis, ancylostoma, toxocara canis, trichuris vulpis, dipylidium caninum, isospora, cryptosporidium, *Bartonella* spp., *Coxiella burnetti* and *Brucella suis* and sarcocystis [8,14,19,20]. Due to the proximity of the Jahai villages to Thailand, there is a risk of the hunting dogs contracting rabies from stray dogs wandering from neighbouring Thailand, where rabies is endemic. In one study conducted in an Orang Asli village, almost 50% of stool samples were positive for helminths [21]. This is probably due to the uncontrolled population of strays and semi domesticated dogs which eat faeces [21] and live in close proximity to human population compounded with poor levels of hygiene, overcrowding and lack of veterinary attention [22]. In some locations despite the provision of better housing and education, most Orang Asli still do not appreciate the importance of personal hygiene, proper usage of toilets and sanitation [21], they prefer walking barefooted, using river water as a source of bathing and cleaning and they hardly use soap or wash before eating [21], making them at higher risk to dog associated infections [11,23].

Effective methods of controlling dog associated zoonotic infections is vaccination, and by increasing the levels of knowledge regarding dog associated zoonotic diseases, recognition of its clinical signs in both humans and animals, promoting good attitudes and practices of the community like washing hands after direct contact with dogs and dogs’ body secretion such as saliva, urine and faeces, taking proper precautions against dog bites and seeking treatment once bitten by dogs, proper washing of vegetables and cooking of meals. However, the success of any programme depends on the cooperation and participation of the community [23].

There have been several reported successful intervention programmes on zoonotic diseases and neglected tropical diseases [24–26] in communities where illiteracy or low levels of literacy is prevalent. In a pilot project in Chile, a ‘One Health’ approach was used to detect, treat and prevent zoonotic infections and it was found that an integrated multidisciplinary programme was valuable in preventing, diagnosing and treating zoonotic infections [24–27]. One Health is a collaborative effort of multiple disciplines working locally, nationally and globally to attain optimal health in a holistic manner for people, animals and environment [28–30] leading to health improvement and optimisation of risk mitigation simultaneously across all domains [31].

Low levels of education of the Jahai compounded with poor hygiene especially hand hygiene, poor nutrition, without supply of treated water and electricity, toilet facility and the distance from health centres expose this already marginalized population to risks of zoonotic infections from dogs. The remoteness of these villages and the accessibility via a boat has rendered very little opportunity for health education and promotion up to now.

The objective of this study was to instil good knowledge, attitude and practices of the Jahai indigenous people in Malaysia concerning dog associated zoonotic infections using One Health approach.

## Methodology

***Study design***: this is a non-experimental pre and post-test intervention study using One Health approach. ***Setting***: This study was carried out in the Jahai Orang Asli settlement in Sungai Tiang village located in the Royal Belum Forest in Perak, one of the 14 states in Malaysia. There are approximately 437 villagers making up 102 families in this village. However because some of them still practice nomadic lifestyles, the actual number of villagers fluctuate from time to time. ***Sampling***: All Jahai villagers aged 12 years and above were eligible to participate in this study. Using Stata to calculate sample size, a sample size of 170 would have been required for the study to have a ± 3% acceptable margin of error. However it was the intention of the investigators to recruit approximate 180 respondents to ensure validity of the findings. The respondents were chosen using a convenient sampling method considering that the population would fluctuate either because some villagers were still nomadic or because most of the villagers would be out either hunting or foraging for food in the forests usually for several days. Each household was approached by the investigators who would invite everyone aged above 12 to participate. ***Tools***: A questionnaire specially designed for this study was used. The interviews were conducted in the participant’s homes using face to face interview by trained investigators using a uniform protocol which was set up to minimize error and bias. Besides the baseline demographic information which consists of sex, age, race, marital status, education level and employment status, information was collected on the knowledge, attitudes and practices of the participants towards dog associated zoonotic infections. The questions on knowledge included if the participants knew about dog associated infections and if it can transmit from stray and pet dogs, how infections are transmitted, route and types as well as the prevention, treatment and detection. The Cronbach’s Alpha for the 42 item knowledge scale for pre and post intervention was 0.73 and 0.72 respectively. Questions on attitude included importance of washing before eating and touching dogs, if they would seek treatment if bitten by a dog and if they were concerned about dog associated infections and were interested to know more. The Cronbach’s Alpha for the 6 item attitude scale for pre and post intervention was 0.71 and 0.75 respectively. Questions on practice included hand washing practices and how they would treat a suspected dog associated infection. The Cronbach’s Alpha for the 9 item practice scale for pre and post intervention was 0.57 and 0.47 respectively. The first part of the study commenced in April 2017 with the preliminary data collection followed by four episodes of health education and promotion relating to dog associated infections. The intervention involved two teams which included medical, veterinary and allied health students. The health intervention materials were jointly developed by the medical and veterinary professionals using One Health concept. ***Intervention***: Health education intervention was done using a face to face approach. Each team of three or four visited each household and basic education on dog behaviour, wellness program and zoonosis transmission from dog to human was conducted using educational posters, slide presentations, comics and video clips with relevant content and messages. This approach has been proven to be highly successful in increasing the attentiveness of both children and adult, leading to better understanding of basic concepts on transmission and preventive measures of zoonotic diseases [26]. Some local terms were used during the intervention programme to enhance comprehension of the discussed topic to ensure the participants feel comfortable to engage with the investigators. Using games, toys and video games, the children were taught on hand washing and dog associated zoonotic infections. They were also taught on proper hand washing techniques. Considering the low levels of education among the Orang Asli, the intervention mostly included pictorial images on posters and flyers which were left with the households. The participants of each household were told to share the information they learned with other household members who were not present. One intervention session was held in the school, after obtaining the permission of the school headmaster, because it has been shown that the access and receptivity level is higher in schools. The villagers were also given a hand-out bag containing soap, tea, sugar and coffee and sandal to ensure participation [25]. The investigators were trained on the subject matter and interpersonal and communication skills to ensure that they were able to connect with the villagers effectively [25]. A post intervention survey was conducted five months later to determine the level of success of the project. ***Analysis***: Data was analysed using PASW version 18. The proportion of each answer was compared using a chi square test for statistical significance. The pre and post mean knowledge, attitudes and practices scores were analysed using t test. A probability value of p<0.05 was considered to be significant. ***Ethics***: The research was conducted ethically. All respondents were read the participants information sheet which contained the information concerning the study including the rights of the participant to refuse and exit from the study at any moment. Each participant was asked for an informed verbal consent before proceeding with the questionnaire. The study received Institutional Ethics Committee (PMC RC8) approval. The anonymity of the respondents is assured.

## Results

As shown in table 1, most of the participants in the pre intervention group were aged 21 and below (45.2%), female (50.3%), animist (76.8%), married (62.7%), highest level of education up to secondary school (41.2%) and most of the households had average monthly household income of RM870 (USD 1= RM 4) and below (88.7%). Most of the participants in the post intervention group were aged 21 and below (46.6%), male (53.3%), animist (70.3%), married (63.7%), with the highest level of education up to secondary school (39.6%), and most of the households had an average monthly household income of RM870 and below (96.7%)

**Table 1:** Demographics of the participants

| **Variables** | **Pre (N= 177)** | **Post (N=182)** |
| --- | --- | --- |
| **A. Demographics** | **n (%)** | **n (%)** |
| **Age** |  |  |
| ≤21 | 80 (45.2) | 85 (46.7) |
| 22-40 | 74 (41.8) | 74 (40.7) |
| 41-59 | 21 (11.9) | 16 (8.8) |
| ≥60 | 2 (1.1) | 7 (3.8) |
| **Sex** |  |  |
| Male | 88 (49.7) | 97 (53.3) |
| Female | 89 (50.3) | 85 (46.7) |
| **Religion** |  |  |
| Animist | 136 (76.8) | 128 (70.3) |
| Islam | 37 (20.9) | 52 (28.6) |
| Christian | 4 (2.3) | 2 (1.1) |
| **Marital status** |  |  |
| Married | 111 (62.7) | 116 (63.7) |
| Divorced / widow | 1 (0.6) | 0 (0.0) |
| Widow | 7 (4.0) | 6 (3.3) |
| Single | 58 (32.8) | 60 (33.0) |
| **Highest Level of Education** |  |  |
| Illiterate | 35 (19.8) | 39 (21.4) |
| Informal | 12 (6.8) | 17 (9.3) |
| Primary | 57 (32.2) | 54 (29.7) |
| Secondary | 73 (41.2) | 72 (39.6) |
| **Monthly Household Income** |  |  |
| ≤870 | 157 (88.7) | 176 (96.7) |
| >870 | 20 (11.3) | 6 (3.3) |

As shown in figure 1, although in general most aspects of the knowledge improved positively post education, there were still a substantial proportion of the respondents who still believed that dog associated infections are spread by smell, aura, by looking at a dog, and that the route of infection was through spirits. Also substantial proportion believed that traditional rituals and prayers can prevent dog associated infections, and dreams and intuition and shaman can help detect infections. The increase in the knowledge post intervention that infections can spread from pet dogs was statistically significant (X^2^=4.293, p= 0.038)

**Figure 1:**
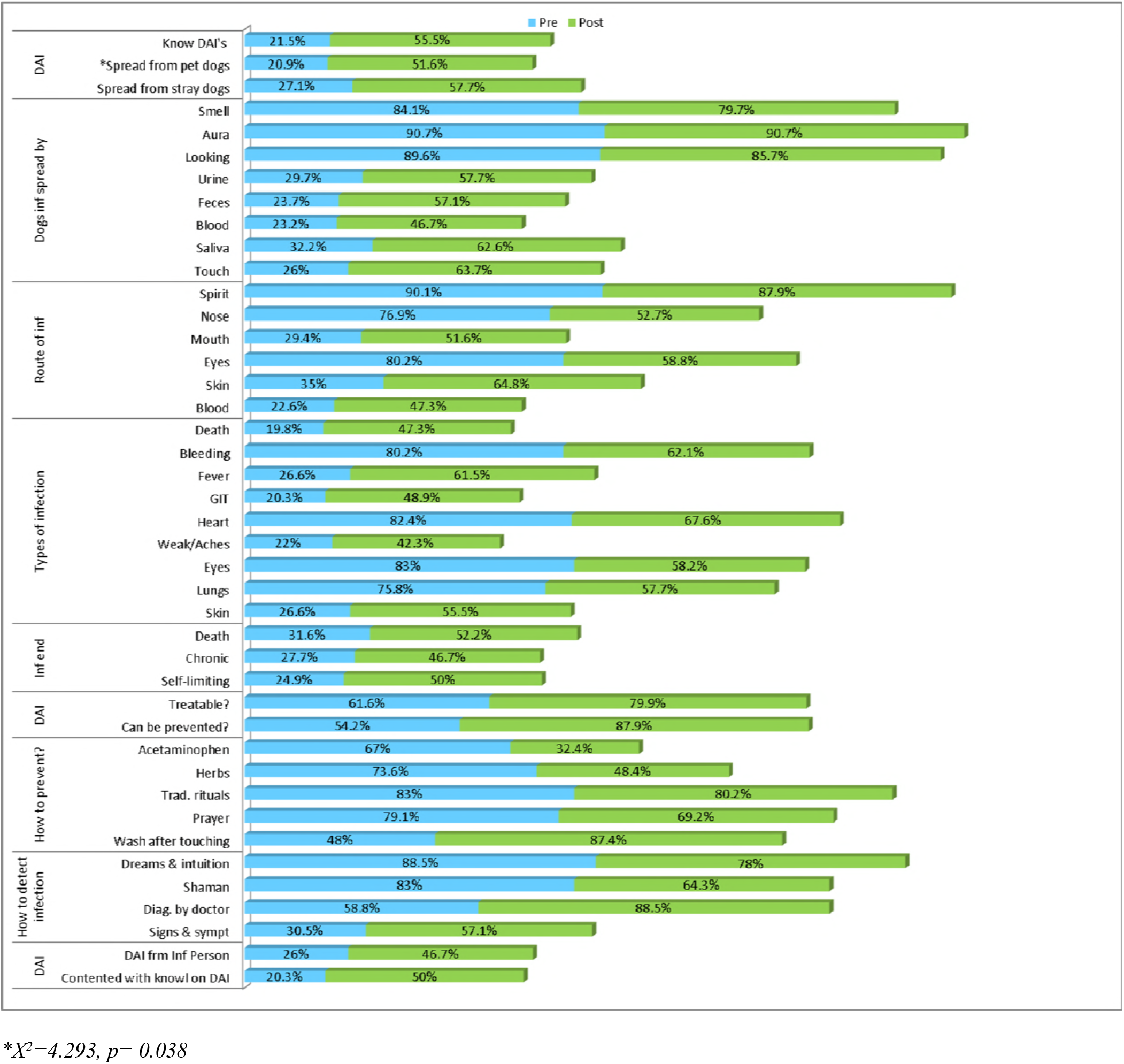
pre and post intervention knowledge concerning dog associated illnesses

In general, there was improvement in all aspects of attitude post intervention (figure 2).

**Figure 2:**
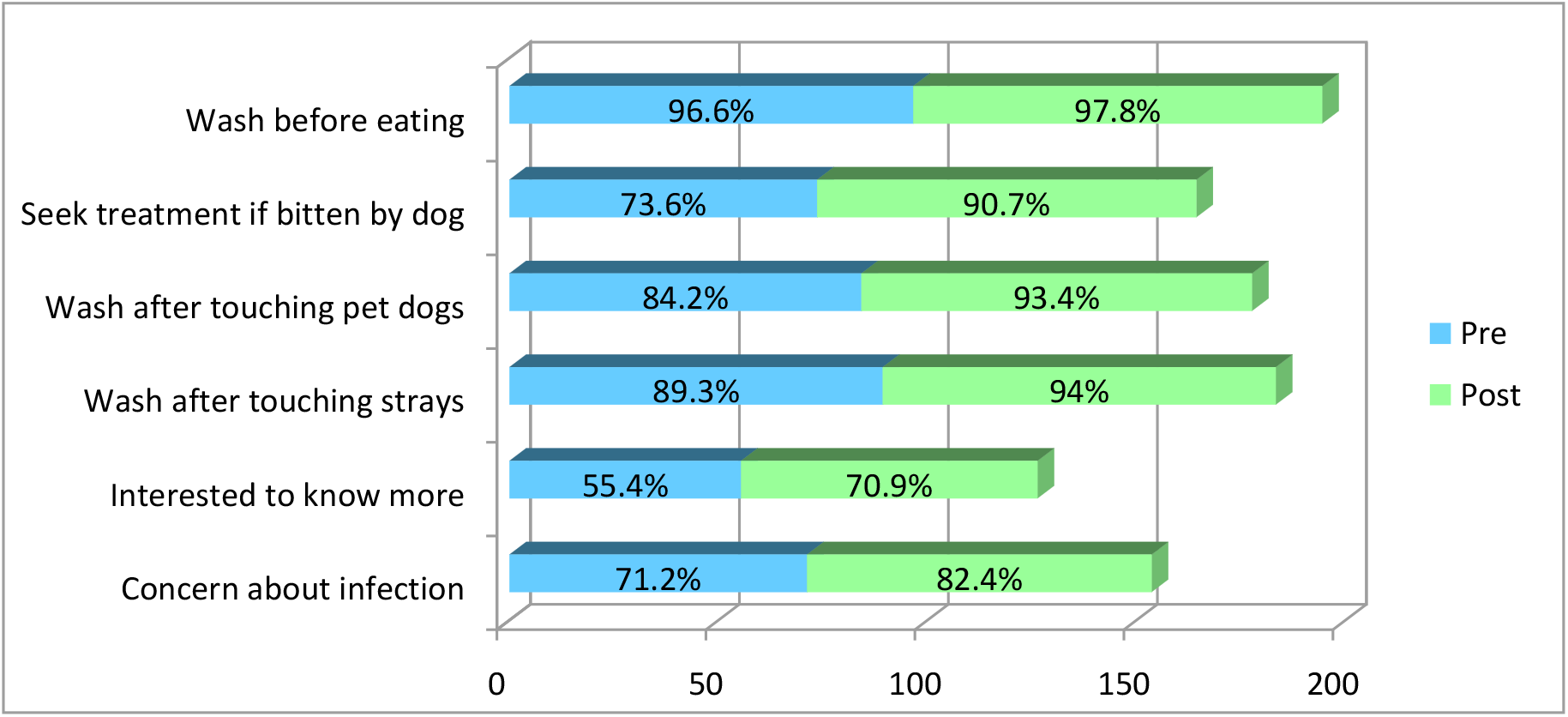
pre and post intervention attitude related to dog associated infections

In general most aspects of practice improved positively post education, however a substantial proportion still believed that dog associated infections can be treated by prayers and by seeking treatment from a shaman. The practice of washing hands before eating improves significantly (X^2^=14.984, p <0.001) post intervention (figure 3).

**Figure 3:**
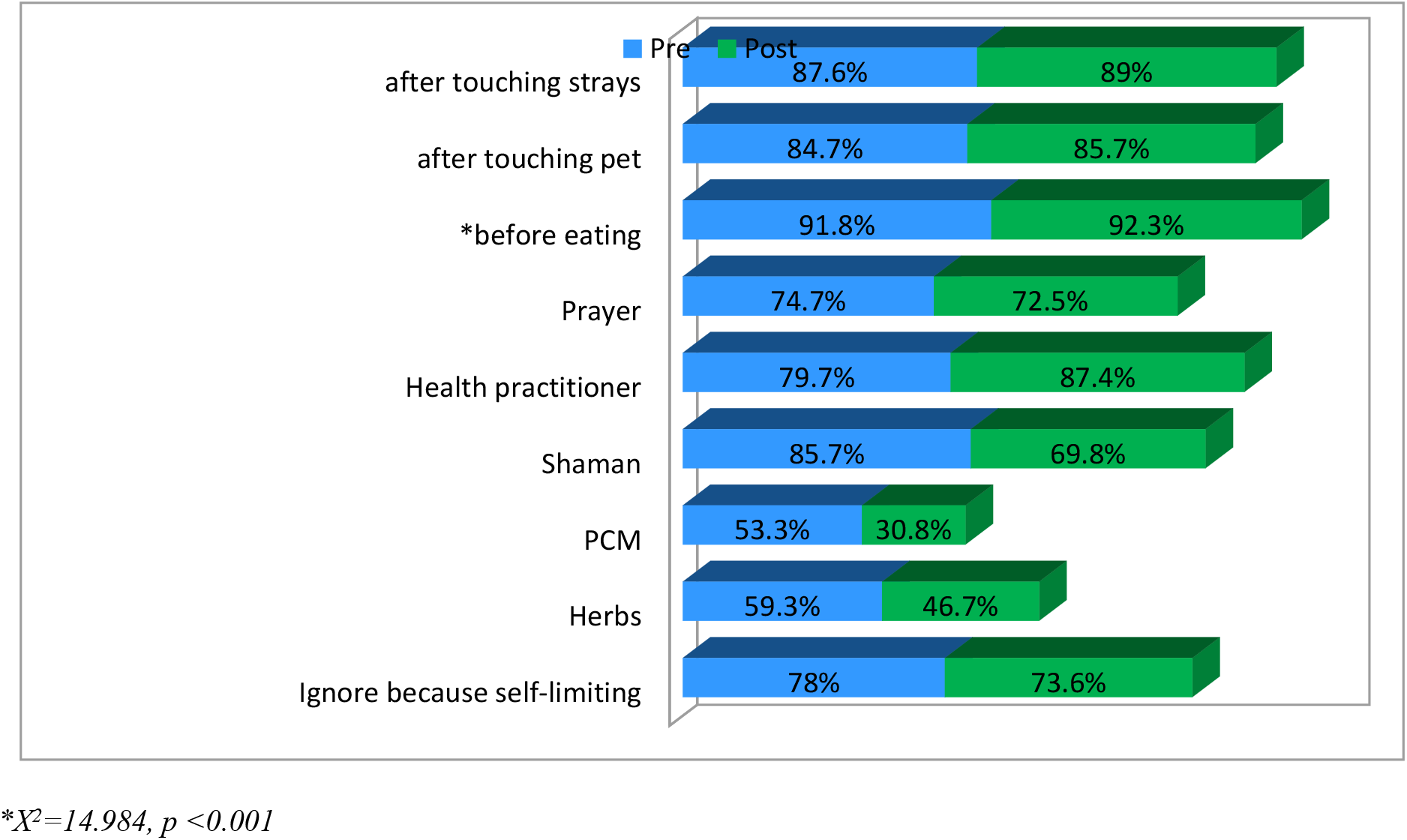
pre and post intervention practices related to dog associated infections

As shown in table 2, the mean score of the participants knowledge increased from 19.3 to 24.7 (t=−9.875, p=<0.001) and the mean attitude score increased from 4.7 to 5.3 (t= −4.100, p=<0.001). The improved mean scores for both knowledge and attitude were statistically significant. There was a slight increase in the mean score for practice before (6.3) and after (6.4) intervention. However this increase was not statistically significant (t=0.057, p=0.940).

**Table 2:**
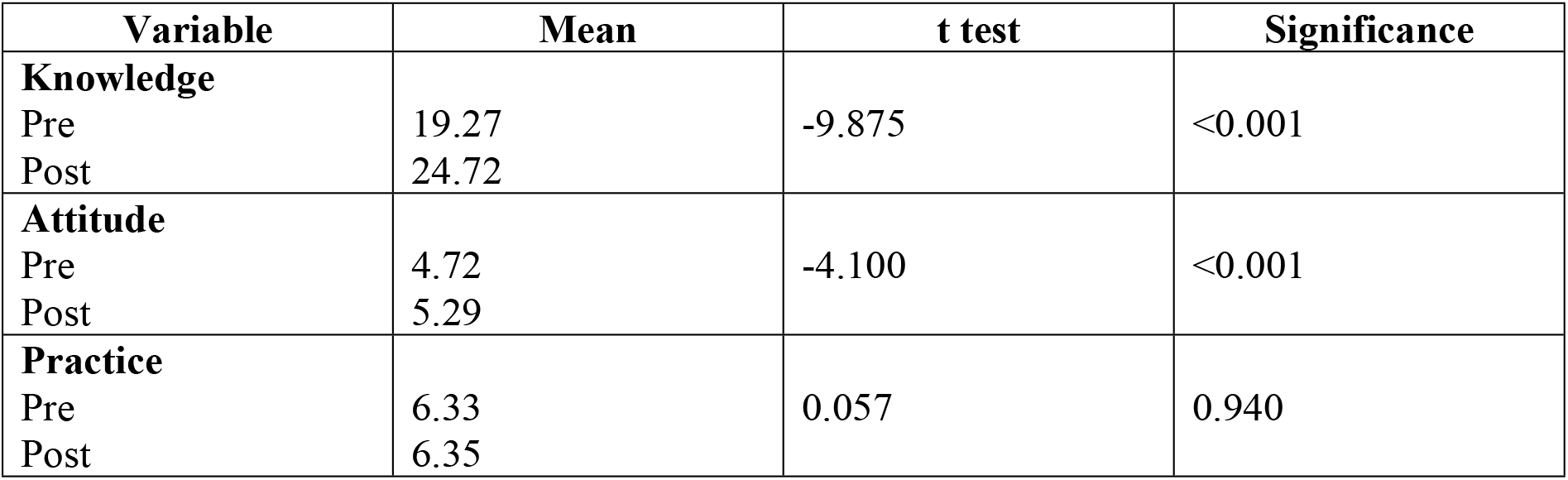
pre and post intervention mean score comparison

## Discussion

### Knowledge

Interventions in forms of training, education and health promotion can increase knowledge of the community [32]. The findings of the current study showed an increase in the knowledge of Orang Asli on dog-led zoonotic infections following health education and promotion. Similarly an evaluation of rabies prevention and control project among dog owners in Philippines showed a significant increase in the knowledge of the community concerning rabies [33]. A classroom-based lesson on rabies among primary school children in Malawi also showed an increased in the knowledge of the children on rabies and how to be safe around dogs [34]. Increasing the knowledge of zoonotic infections is important and can reduce dog associated zoonotic infections exponentially [14] but low educational levels, poor access to information, distance of health facilities [35], poverty, lack of treated water and proper faecal disposal in the Orang Asli communities [21] add to the disinterest concerning dog associated illnesses and are impediments to a successful prevention programme [36]. In India, most of the fatalities related to rabies occured due to ignorance and lack of accessibility to affordable services [36]. In Brazil it is reported that lack of knowledge is a reason for the high risk of canine intestinal parasites [19]. Some authors have attributed poor knowledge to medical, allied health professionals and veterinarians who do not discuss dog associated diseases with the public [11,13,35–39].

In general, the knowledge of dog associated infections is low in most developing countries except for fatal infections like rabies which receive a lot more publicity compared to other dog associated illnesses. Although the finding of the present study showed improvement in the knowledge of the villagers post intervention but it is still lower than studies conducted among dog owners in high prevalent rabies locations like Ethiopia, Nigeria and India. But even in these countries, in most cases this information is restricted to rabies, the knowledge concerning parasitic infections and their modes of transmission is, however, limited [18,35,36,40,41]. This is not surprising considering people are generally more concerned about illnesses which receive more media attention, for example because human hydatid cyst is common in Tibet hence the awareness of cystic echinococcosis is fairly high in communities there [23]. Considering knowledge is dependent on the level of education [40,41], it is not surprising the knowledge of the Orang Asli in this study is poor compared to studies conducted in developed nations like the United States of America and Australia where in some cases the participants were even aware of the signs of infections like leptospirosis, giardiasis, rabies, hookworm, coccidiosis, lyme disease, roundworms, toxoplasma, leishmaniais, salmonellosis, ringworm, hookworm, hydatids, toxocaris, giardiasis, toxoplasmosis [11,13,37,38]. The Orang Asli community in this study have no access to information via the internet and other sources due to no connectively at all which also contributes to the low levels of knowledge concerning zoonotic infections [14,42].

### Attitude

The attitude to dog associated infections was good and improved post intervention but further improvement is dependent on the knowledge of the community because lack of basic knowledge in diseases transmission has direct influence on positive attitudes and practices [39]. In general, studies conducted in Philippines [33], China [23], Nigeria [40], Canada [39] and Australia [13] showed good attitudes towards dog associated infections. In Nigeria people felt that children should not play with unknown dogs and that they would wash wound after a dog bite with soap and water [35,40].

Studies have showed that absence of diseases like rabies in some countries could be a reason why people are not very concerned about catching diseases from pets [13,39], are less likely to seek treatment if bitten by a dog [13] and are happy with the level of information they have [39].

### Practice

The findings of the present study showed no significant increase in practice post-test. The reason for this could be because it takes longer for knowledge to be translated into practice as behaviours take longer time to change. Hand washing with soap after handling animals and cleaning the bite wound with plenty of running water is important in controlling zoonotic infections. Simple measures such as correct hand washing technique is an effective and a low-cost intervention that is able to provide great health benefit particularly in rural or aboriginal areas. Hand hygiene serves a critical role in reducing the risk of zoonotic infections and is a key measure in preventing dog associated zoonotic infections in the community [13,14,43]. Because of the cost effectiveness, hand washing is often emphasised in Orang Asli schools and communities. This could explain the high prevalence of the practice even before intervention in this study. In a study in Tibet China most respondents always often wash their hands before eating even when other practices on disease prevention and control was poor [23]. Studies in Canada showed high prevalence of hand washing among pet owners and their children after touching pets [14].

Unavailability of treated water in the Orang Asli community may be an impediment to hand hygiene practices because besides knowledge, the effectiveness of hand hygiene practice is dependent on many other factors including socio-economic status, culture and most importantly the availability of clean water and soap [41,44]. For example in Ethiopia although pet owners agreed that washing hands with soap after coming in contact with pets is a preventive measure but the proportion that washed was small [41].

Discarding old beliefs is difficult and will take time. In Nigeria people believed that rabies can be transmitted by air and that it can be treated with traditional medicines [35]. In India wound management of dog bites included oils, turmeric powder and red chillies [45] and although a big majority would consult a doctor for dog bites but substantial number of people preferred using herbal medicines [36]. Even when wounds were cleaned, the methods of cleaning were not be effective in disease prevention [13].

## Conclusion

Both veterinarians and health practitioners have an important role to play in the prevention of dog associated zoonotic infections by educating the public. Although there were significant improvements in the level of knowledge and attitudes of the community before and after intervention, it is difficult to measure the exact and precise effectiveness of the interventions as different communities have different behaviours and require different methods of evaluation and benchmarks [46,47]. The authors suggest that a sustained commitment to a continuous health education and promotion interventions involving the community and school children which promote knowledge and sharing of skills be custom made for indigenous communities [14,25,26,38,48–50] and sanitation and hygienic practices are important areas that must be reinforced at every opportunity [51,52]. These Orang Asli villages provide opportunities for multidisciplinary professionals to work together using One Health approach to provide health education and promotion concerning zoonotic infections and can be a good training ground for future one health workforce.

## Acknowledgement

The authors would like to thank USAID and Malaysia One Health University Network, without whom the study would not have been a success.

## References

1. Affairs TDoEaS (2009) The state of the worlds indigenous people. New York: United Nations.

2. Gomes AG (2004) The Orang Asli of Malaysia. International Institute for Asian Studies Newsletter 35.

3. Ali ZAH (2003) Pentadbiran Orang asli: kemana hala tujunya pengalaman dan cabaran. In: Department of Museum and Antiquities and the Ministry of Art CaHM, editor. warisan Orang Asli Kuala Lumpur: Muzium Negara.

4. Gomes AG (2007) Modernity and Malaysia: settling the Menraq forest nomads. London and New York: Taylor & Francis. 200 p.

5. Schwabe KA, Carson RT, DeShazo JR, Potts MD, Reese AN, et al. (2014) Creation of Malaysia’s Royal Belum State Park: A Case Study of Conservation in a Developing Country. The Journal of Environment & Development 24: 54–81.

6. Kasim Z, Baskaran D (2014) The life of indigenous peoples of Belum-Temonggor: Yayasan Emkay. 197 p.

7. Gianno R, Bayr KJ (2009) Semelai Agricultural Patterns: Toward an Understanding of Variation among Indigenous Cultures in Southern Peninsular Malaysia. Journal of Southeast Asian Studies 40: 153–185.

8. Ghasemzadeh I, Namazi SH (2015) Review of bacterial and viral zoonotic infections transmitted by dogs. Journal of Medicine and Life 8: 1–5.

9. Lee AC, Schantz PM, Kazacos KR, Montgomery SP, Bowman DD (2010) Epidemiologic and zoonotic aspects of ascarid infections in dogs and cats. Trends Parasitol 26: 155–161.

10. Oehler RL, Velez AP, Mizrachi M, Lamarche J, Gompf S (2009) Bite-related and septic syndromes caused by cats and dogs. Lancet Infect Dis 9: 439–447.

11. Bingham GM, Budke CM, Slater MR (2010) Knowledge and perceptions of dog-associated zoonoses: Brazos County, Texas, USA. Preventive Veterinary Medicine 93: 211–221.

12. Patronek GJ, Slavinski SA (2009) Animal bites. J Am Vet Med Assoc 234: 336–345.

13. Steele SG, Mor SM (2015) Client knowledge, attitudes and practices regarding zoonoses: a metropolitan experience. Australian Veterinary Journal 93: 439–444.

14. Stull JW, Peregrine AS, Sargeant JM, Weese JS (2013) Pet husbandry and infection control practices related to zoonotic disease risks in Ontario, Canada. BMC Public Health 13: 520.

15. Mani I, Maguire JH (2009) Small animal zoonoses and immuncompromised pet owners. Top Companion Anim Med 24: 164–174.

16. Stull J, Stevenson K (2015) Zoonotic disease risks for immunocompromised and other high-risk clients and staff-promoting safe pet ownership and contact. Veterinary Clinics of North America: Small Animal Practice 45: 377–392.

17. Thompson RC (2000) Giardiasis as a re-emerging infectious disease and its zoonotic potential. Int J Parasitol 30: 1259–1267.

18. Ugbomoiko US, Ariza L, Heukelbach J (2008) Parasites of importance for human health in Nigerian dogs: high prevalence and limited knowledge of pet owners. BMC Vet Res 4: 1746–6148.

19. Katagiri S, Oliveira-Sequeira TC (2008) Prevalence of dog intestinal parasites and risk perception of zoonotic infection by dog owners in Sao Paulo State, Brazil. Zoonoses Public Health 55: 406–413.

20. Shapiro AJ, Brown G, Norris JM, Bosward KL, Marriot DJ, et al. (2017) Vector-borne and zoonotic diseases of dogs in North-west New South Wales and the Northern Territory, Australia. BMC Veterinary Research 13: 238.

21. Sinniah B, Sabaridah I, Soe MM, Sabitha P, Awang IP, et al. (2012) Determining the prevalence of intestinal parasites in three Orang Asli (Aborigines) communities in Perak, Malaysia. Trop Biomed 29: 200–206.

22. Traub RJ, Robertson ID, Irwin PJ, Mencke N, Thompson RC (2005) Canine gastrointestinal parasitic zoonoses in India. Trends Parasitol 21: 42–48.

23. Li D, Gao Q, Liu J, Feng Y, Ning W, et al. (2015) Knowledge, attitude, and practices (KAP) and risk factors analysis related to cystic echinococcosis among residents in Tibetan communities, Xiahe County, Gansu Province, China. Acta Tropica 147: 17–22.

24. Bardosh K, Inthavong P, Xayaheuang S, Okello AL (2014) Controlling parasites, understanding practices: the biosocial complexity of a One Health intervention for neglected zoonotic helminths in northern Lao PDR. Soc Sci Med 120: 215–223.

25. Ducrotoy MJ, Yahyaoui Azami H, El Berbri I, Bouslikhane M, Fassi Fihri O, et al. (2015) Integrated health messaging for multiple neglected zoonoses: Approaches, challenges and opportunities in Morocco. Acta Trop 152: 17–25.

26. WindaWidyastuti MD, Jatikusumah A, Sunandar, Basuno E, Putra AAG, et al. (2014) Village Rabies Working Group: The Knowledge Translation of Understanding Better on Dog Ecology and Community Engagement for Optimizing Rabies Control Program in Bali, Indonesia. Ecohealth Conference. 2014.

27. Peña A, Abarca K, Weitzel T, Gallegos J, Cerda J, et al. (2016) One Health in Practice: A Pilot Project for Integrated Care of Zoonotic Infections in Immunocompromised Children and Their Pets in Chile. Zoonoses Public Health 63: 403–409.

28. Evans BR, Leighton FA (2014) A history of One Health. Rev Sci Tech 33: 413–420.

29. Gibbs EPJ (2014) The evolution of One Health: a decade of progress and challenges for the future. Veterinary Record 174: 85.

30. King LJ, Anderson LR, Blackmore CG, Blackwell MJ, Lautner EA, et al. (2008) Executive summary of the AVMA One Health Initiative Task Force report. J Am Vet Med Assoc 233: 259–261.

31. Lebov J, Grieger K, Womack D, Zaccaro D, Whitehead N, et al. (2017) A framework for One Health research. One Health 3: 44–50.

32. Olalekan AW, Adebukola AM (2015) Effects of Training on Knowledge, Attitude and Practices of Malaria Prevention and Control among Community Role Model Care Givers in South Western Nigeria. Ethiop J Health Sci 25: 329–336.

33. Barroga T, Basitan I, Lobete T, Bernales R, Gordoncillo M, et al. (2018) Community Awareness on Rabies Prevention and Control in Bicol, Philippines: Pre- and Post-Project Implementation. Tropical Medicine and Infectious Disease 3: 16.

34. Burdon Bailey JL, Gamble L, Gibson AD, Bronsvoort BMD, Handel IG, et al. (2018) A rabies lesson improves rabies knowledge amongst primary school children in Zomba, Malawi. PLoS Negl Trop Dis 12.

35. Kiflu B, Abdurahaman M, Alemayehu H, Eguale T (2016) Investigation on public knowledge, attitude and practices related to pet management and zoonotic canine diseases in Addis Ababa, Ethiopia. Ethiopian Veterinary Journal 20: 67.

36. B K, S P, A D, N R, G Y, et al. (2016) Knowledge, Attitude And Practices Related To Animal Bites Among The Residents Of An Urbanized Village In South Delhi. International Journal of Research and Development in Pharmacy and Life Sciences 5: 2164–2168.

37. Fontaine RE, Schantz PM (1989) Pet Ownership and Knowledge of Zoonotic Diseases in De Kalb County, Georgia. Anthrozoös 3: 45–49.

38. Sandhu GK, Singh D (2014) Level of Awareness Regarding Some Zoonotic Diseases, Among Dog Owners of Ithaca, New York. Journal of Family Medicine and Primary Care 3: 418–423.

39. Stull JW, Peregrine AS, Sargeant JM, Weese JS (2012) Household knowledge, attitudes and practices related to pet contact and associated zoonoses in Ontario, Canada. BMC Public Health 12: 553–553.

40. Ameh VO, Dzikwi AA, Umoh JU (2014) Assessment of Knowledge, Attitude and Practice of Dog Owners to Canine Rabies in Wukari Metropolis, Taraba State Nigeria. Global Journal of Health Science 6: 226–240.

41. Tensa WF (2017) Household Knowledge, Attitudes and Practices Related to Pet Contact and Associated Zoonosis in Bishoftu, Ethiopia. Global Veterinaria 18: 277–285.

42. Otranto D, Dantas-Torres F, Mihalca AD, Traub RJ, Lappin M, et al. (2017) Zoonotic Parasites of Sheltered and Stray Dogs in the Era of the Global Economic and Political Crisis. Trends Parasitol 33: 813–825.

43. Bloomfield SF, Aiello AE, Cookson B, O’Boyle C, Larson EL (2007) The effectiveness of hand hygiene procedures in reducing the risks of infections in home and community settings including handwashing and alcohol-based hand sanitizers. American Journal of Infection Control 35: S27–S64.

44. McDonald E, Cunningham T, Slavin N (2015) Evaluating a handwashing with soap program in Australian remote Aboriginal communities: a pre and post intervention study design. BMC Public Health 15: 1188.

45. AS S, A S, P K (2002) Micro conceptions and myths in the management of animal bite case. Indian J Community Med 27: 9–11.

46. McLeroy KR, Norton BL, Kegler MC, Burdine JN, Sumaya CV (2003) Community-Based Interventions. American Journal of Public Health 93: 529–533.

47. Santiago AM, Soska TM, Gutierrez LM (2017) Improving the Effectiveness of Community-Based Interventions: Recent Lessons from Community Practice. Journal of Community Practice 25: 139–142.

48. Catalani C, Minkler M (2010) Photovoice: a review of the literature in health and public health. Health Educ Behav 37: 424–451.

49. Darlington EJ, Violon N, Jourdan D (2018) Implementation of health promotion programmes in schools: an approach to understand the influence of contextual factors on the process? BMC Public Health 18: 163.

50. Ubert T, Forberger S, Gansefort D, Zeeb H, Brand T (2017) Community Capacity Building for Physical Activity Promotion among Older Adults-A Literature Review. Int J Environ Res Public Health 14.

51. Bulled N, Poppe K, Ramatsisti K, Sitsula L, Winegar G, et al. (2017) Assessing the environmental context of hand washing among school children in Limpopo, South Africa. Water Int 42: 568–584.

52. Clasen T, Schmidt W-P, Rabie T, Roberts I, Cairncross S (2007) Interventions to improve water quality for preventing diarrhoea: systematic review and meta-analysis. Bmj 334: 782.

